# Unveiling genetic anchors in Saccharomyces cerevisiae: QTL mapping identifies IRA2 as a key player in ethanol tolerance and beyond

**DOI:** 10.1101/2024.05.08.593158

**Authors:** Larissa Escalfi Tristão, Lara Isensee Saboya de Sousa, Beatriz de Oliveira Vargas, Juliana José, Marcelo Falsarella Carazzolle, Eduardo Menoti Silva, Juliana Pimentel Galhardo, Gonçalo Amarante Guimarães Pereira, Fellipe da Silveira Bezerra de Mello

## Abstract

Ethanol stress in *Saccharomyces cerevisiae* is a well-studied phenomenon, but pinpointing specific genes or polymorphisms governing ethanol tolerance remains a subject of ongoing debate. Naturally found in sugar-rich environments, this yeast has evolved to withstand high ethanol concentrations, primarily produced during fermentation in the presence of suitable oxygen or sugar levels. Originally a defense mechanism against competing microorganisms, yeast-produced ethanol is now a cornerstone of brewing and bioethanol industries, where customized yeasts require high ethanol resistance for economic viability. However, yeast strains exhibit varying degrees of ethanol tolerance, ranging from 8% to 20%, making the genetic architecture of this trait complex and challenging to decipher. In this study, we introduce a novel QTL mapping pipeline to investigate the genetic markers underlying ethanol tolerance in an industrial bioethanol *S. cerevisiae* strain. By calculating missense mutation frequency in a prominent QTL region within a population of 1011 *S. cerevisiae* strains, we uncovered rare occurrences in gene *IRA2*. Following molecular validation, we confirmed the significant contribution of this gene to ethanol tolerance, particularly in concentrations exceeding 12% of ethanol. *IRA2* pivotal role in stress tolerance due to its participation in the Ras-cAMP pathway was further supported by its involvement in other tolerance responses, including thermotolerance, low pH tolerance, and resistance to acetic acid. Understanding the genetic basis of ethanol stress in *S. cerevisiae* holds promise for developing robust yeast strains tailored for industrial applications.

## INTRODUCTION

Ethanol stress in yeast, characterized by the adverse effects of cellular exposure to elevated ethanol concentrations, leads to compromised metabolic functionality and diminished cellular viability. A diverse array of physiological and biochemical responses is triggered to mitigate these effects, including alterations in cell membrane fluidity and protein denaturation (Jhariya et al., 2021). While different yeast strains exhibit varying degrees of tolerance to ethanol exposure, *Saccharomyces cerevisiae*, specialized in ethanol production, demonstrates heightened tolerance, offering an evolutionary advantage in sugar-rich environments (Jhariya et al., 2021). As the primary metabolite of *S. cerevisiae*, ethanol can be produced and accumulated at high concentrations by these cells. Over centuries, *S. cerevisiae* has been extensively utilized for food and beverage fermentation, and more recently, for biofuel production (Parapouli et al., 2020).

Bioethanol plays a pivotal role in the global energy landscape as a renewable and sustainable fuel source, with its production being a significant application of *S. cerevisiae*, which can subject the cell to heightened ethanol stress. The fermentation of sugarcane and corn can yield ethanol concentrations of up to 12% and 18% (v/v), respectively (Lopes et al., 2016), due to the efficient conversion of sugars into ethanol through the action of enzymes such as zymase and alcohol dehydrogenase. Despite the numerous adverse conditions that yeast must endure to survive during fermentation processes - including acidic environments, exposure to high temperatures, and inhibitory molecules, such as acetic acid and furfural -, ethanol stress is widely recognized as one of the most significant and challenging limiting factors in industrial bioethanol production (Varize et al., 2022). Even though *S. cerevisiae* exhibits higher ethanol tolerance than other yeasts, this phenomenon compromises cell viability, thereby disrupting fermentation performance and ultimately diminishing ethanol yield (Sahana et al., 2024).

Numerous mechanisms have been identified in *S. cerevisiae* to confer ethanol tolerance. These mechanisms involve the regulation of multiple genes and transcription factors, along with the activation of stress response elements, including heat shock proteins (HSP), osmotic regulators, and oxidative enzymes. Essential for enhancing membrane integrity and restoring membrane fluidity, reorganization, and remodeling of the plasma membrane involves the increased presence of specific compounds such as ergosterol and unsaturated fatty acids. Accumulation of proline and tryptophan plays a vital role in stabilizing proteins and membranes, while induced HSP ensures proper protein folding and stability. The trehalose pathway, in conjunction with HSP, contributes to protein stabilization and alleviates plasma membrane permeabilization. Diverse HSP, including HSP30, HSP32p, HSDP31p, and HSP150, perform various functions essential for ethanol tolerance. Transcription factors such as Yap1p, Msn2p/Msn4p, Hog1, and Hsf1p, along with the ASR1 gene, are implicated in orchestrating this stress response (Jhariya et al., 2021).

Despite extensive knowledge of the response to ethanol stress in *S. cerevisiae*, ethanol tolerance remains a polygenic and complex phenotype involving over 250 genes, which has yet to be fully elucidated (Basso et al., 2011; Jhariya et al., 2021). Moreover, different strains exhibit varying degrees of ethanol tolerance, ranging from 8% to 20%, underscoring the importance of understanding these responses to enhance fermentation performance and develop robust yeast strains for industrial applications. In this regard, quantitative trait loci (QTL) mapping emerges as an efficient technique for uncovering gene mutations that confer higher tolerances, as well as identifying novel phenotype-associated genes (Deutschbauer and Davis, 2005). QTL mapping of industrially relevant traits in industrial strains has proven highly promising, as these strains represent a valuable resource for understanding and exploring intricate metabolism and uncovering gene mutations that confer tolerance, such as resistance to acidic pH, thermotolerance, and aldehydes (Coradini et al., 2021; de Mello et al., 2022a; Ho et al., 2021; Swinnen et al., 2012; Wang et al., 2019).

In this study, we delved into the mechanisms underpinning the high ethanol tolerance trait of the Brazilian industrial strain SA-1 using a QTL mapping approach. We developed a novel QTL discovery pipeline tailored for swiftly identifying alleles and mutations associated with complex phenotypes. This effort led to the identification of a key mechanism linked to ethanol tolerance. Furthermore, this allele displayed correlations with other tolerance mechanisms that yeast encounter during fermentation processes, highlighting its significance in coping with environments characterized by multiple stresses.

## MATERIAL AND METHODS

### Strains and growth media

A total of 78 yeast strains (**Table S1**) were used for screening ethanol tolerance in yeast. The main strains used for QTL mapping are described in **Table 1**. The strains were cultivated in YPD medium (20 g.L^-1^ of peptone, 10 g.L^-1^ of yeast extract, and 20 g.L^-1^ glucose) for pre-inoculum or control conditions. For ethanol tolerance screening, YPD supplemented with an ethanol solution at final concentrations of 8 up to 18% (v/v) was used. For other tolerance tests, YPD was supplemented with 8 to 12 g/L of acetic acid with pH adjusted to 4,7; and H_2_SO_4_ to pH of 2.1 or 2.5. Thermotolerance was assessed with cultivation in YPD at 41 °C and 42 °C. All cultivations occurred at 30 °C unless stated otherwise and 250 rpm when agitation was necessary. 15 g.L^-1^ agar was added to YPD for solidification.

**Table 1:** Main *S. cerevisiae* strains used in this study.

### Characterization of ethanol resistance

To evaluate ethanol resistance on solid media, both colony size and serial dilutions were utilized to determine the phenotype. For colony size assessment, an inoculum was prepared in 150 µL YPD in 96-well microplates, and strains were cultured overnight. Subsequently, yeasts were plated on YPD with ethanol plates using a 96-pin replicator and incubated for up to 96 hours. Strain growth was quantified by measuring the area of colonies, as previously outlined (Matsui and Ehrenreich, 2016). The total pixel intensity within a circle (with a dot radius of 50 pixels) around each colony on the image was measured using the Plate Analysis JRU v1 plugin, obtained from the Stowers Institute ImageJ. Colony size was assessed in triplicates. Relative colony size was determined as the ratio between the colony size under ethanol treatment and the control growth conditions. Spot assays were conducted using serial dilutions of a culture with OD_600_ 1.0 and incubated for the required duration.

Phenotyping in liquid media was conducted in 96-well flat-bottom microplates by adding 135 µL YPD supplemented with an ethanol solution and 15 µL inoculum with OD_600_ 1.0 in quadruplicates. The plates were sealed with a MicroAmp film Translucent gas-tight (Applied Biosystems), and growth was assessed by measuring OD_600_ using a microplate reader (SpectraMax Plus 384). The maximum specific growth rate (μ_MAX_) was calculated from the collected data (OD_600_ *vs* Time) using the OCHT software (De Mello et al., 2019). Significant differences in growth parameters were calculated using a *t* test.

### QTL mapping

Mapping of the loci related to the phenotype was performed as previously described (de Mello et al., 2022a). In short, the heterozygous hybrid SA/CEN was sporulated and the haploid segregants were collected manually and further phenotyped in 18% (v/v) ethanol. Two populations of 60 individuals were selected according to their tolerance towards the alcohol, either resistant or susceptible. The total genomic DNA (gDNA) of the two populations was pooled extracted and sequenced using the Illumina HiSeq 4000 platform. The generated 2×100 paired-end reads were initially aligned to the genome sequence of the susceptible parental CEN.PK113-1A, and bulk segregant analysis was performed according to (Takagi et al., 2013). The parameters Gprime, p-value, and Δ(SNP-index) were used to infer the regions of interest.

### SNP frequency in an *S. cerevisiae* population

The frequency of mutations in the gene *IRA2* was identified from the genomic and phenotypic data of 1011 publicly available yeast strains (PETER, 2018) using an in-house-prepared Python script (https://github.com/edumenotti/IC_scripts). Firstly, gene prediction was performed on the 1011 *S. cerevisiae* genomes using the Augustus software (Stanke et al., 2008) with a pre-trained model of *S. cerevisiae RM11-1a*, and the quality of the prediction was assessed using the BUSCO tool (Simão et al., 2015) with Saccharomycetales database. Proteomes with at least 90% completeness of the genomes were clustered into homologous gene families using the OrthoFinder v2.2.6 algorithm (Emms and Kelly, 2019). The proteins from each orthogroup were then globally aligned using MAFFT v7 software (Katoh and Standley, 2013), producing mafft alignment files from which mutation identification and allelic frequency calculations were conducted. Using the in-house Python script, *IRA2* mutations were scanned in the mafft alignment of the orthogroup to which the Ira2p protein belongs, thus calculating the frequency at which the detected mutations occur at those positions compared to all other proteins in the orthogroup. Mutations that have passed a frequency filter of up to 10% - a criterion adopted to consider them rare in comparison with all proteins in the orthogroup - are recorded in a text file, detailing both the mutated amino acid and the frequency at that position.

### Reciprocal hemizygosity analysis

To test the influence of *IRA2* alleles on the stress response, the gene was knocked out individually and further crossed with the non-deleted remaining parent, rendering strains SA (SA49/CEN.PK113-1A *ira2*Δ::hphMX6) and CEN (SA49 *ira2*Δ::hphMX6/CEN.PK113-1A). Genetic engineering was performed with a CRISPR-Cas9 system previously described (de Mello et al., 2022b). Overall, a single-guided RNA (sgRNA) with homology to the *IRA2* open reading frame was used to insert a hygromycin resistance marker (*hphMX6*) in the target region, inactivating the endogenous coding sequence. Yeasts were transformed with the LiAc protocol, null mutants were collected in YPD plates with hygromycin (300 µg/mL), and the edition was confirmed by PCR. sgRNA sequence and primers used are provided in **Table S2**.

## RESULTS

### Selection of an ethanol-tolerant strain

For a QTL mapping approach, assessing the inhibitory concentration for a given stress is essential. First, a parental diploid yeast with a notable phenotype must be selected, followed by the generation of an F_1_ progeny for screening of haploid segregants with traits resembling those of the diploid parental strain. Therefore, 78 yeast strains were screened in increasing ethanol concentrations (8 to 14% v/v) in solid media (**Figure S1**). Laboratory haploid *S. cerevisiae* strains were used as controls. After 96 hours of growth, the relative colony size for all tested concentrations was calculated (**Figure 1A**). All ethanol concentrations hindered yeast growth, even at the lowest concentration tested (8%). However, more ethanol partially restored the performance of some strains. At 14% ethanol, although some strains exhibited significant growth, several yeast strains showed no cells at this concentration after 96 hours. It is also noteworthy that, for all concentrations except 8% ethanol, certain yeast strains exhibited enhanced performance compared to control conditions (relative colony size greater than 1.0).

**Fig 1.**
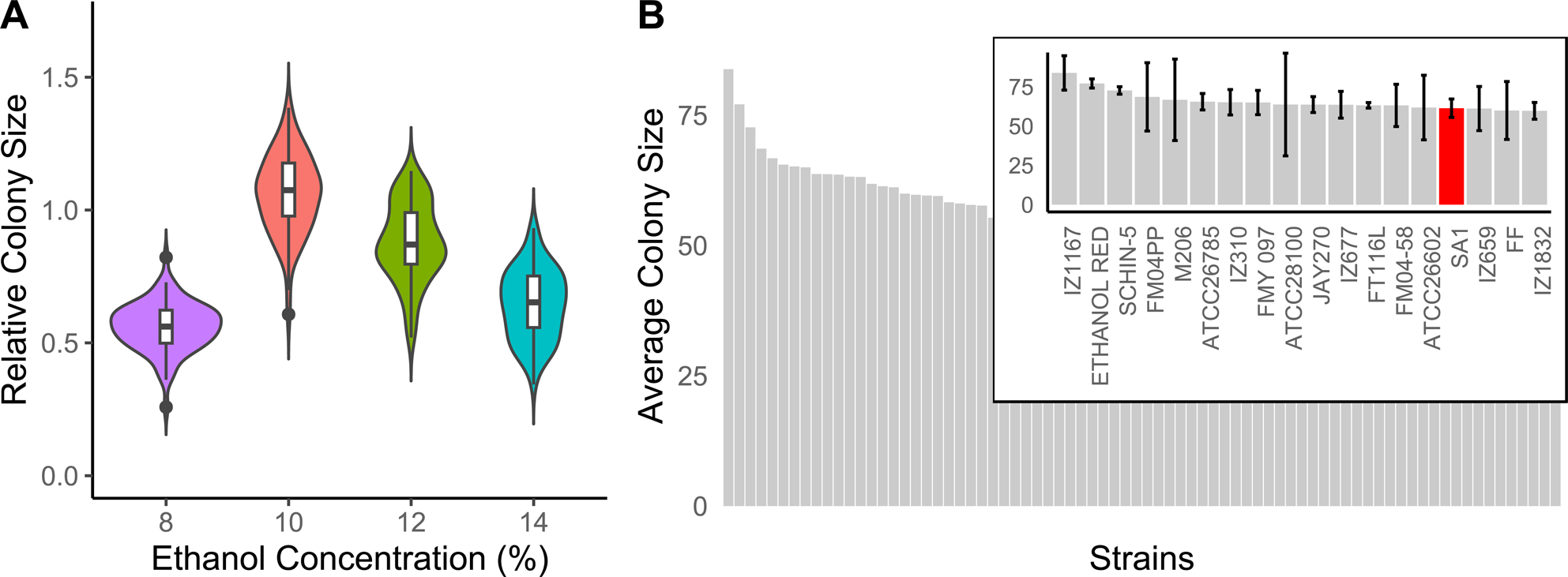
Screening of 78 *Saccharomyes* spp. yeasts in ethanol challenges in solid media at 30 °C. A) Distribution of the relative colony size of each strain for all ethanol concentrations tested. b) Average colony size of all strains at 14% ethanol.

After selecting 14% ethanol as the threshold concentration for collecting ethanol-resistant yeast, colony size at this concentration was evaluated in triplicate (**Figure 1B**). The top 18 strains from this assay are highlighted in the figure. Among these strains, IZ1167 ranked highest under this condition, exhibiting an average colony size of 83.83 ± 10.94. Despite being identified as *S. cerevisiae*, sporulation was not observed after multiple attempts, hindering the QTL mapping pipeline. Following the identification of top-performing yeasts, notable bioethanol industrial strains emerged, including Ethanol Red, JAY270, SA-1, and FMY097 (SA-1 segregant). Given the availability of a segregant library and SA-1’s documented tolerance to various industrial stresses (de Mello et al., 2022a; De Mello et al., 2019), it was selected as the parent strain for this study, displaying an average colony size of 61.37 ± 5.86.

Following the selection of the Brazilian industrial *S. cerevisiae* SA-1 as the background for studying the genetic architecture related to ethanol resistance, a library of 113 segregant haploids was screened under ethanol stress. Initially, a pre-screening in 12% ethanol was conducted to eliminate susceptible strains, resulting in only 22 strains exhibiting growth comparable to the parent in the first screening. Increasing the ethanol concentration to 14% impaired the performance of most strains, consistent with observations from the initial assay with the 78-yeast library (**Figure S2**). At this concentration, haploid SA49 demonstrated the most favorable outcome, with a relative colony size of 0.78. To validate this phenotype, a spot assay was conducted using SA-1, SA49, laboratory strain CENPK113-1A (which performed poorly in the initial screening), and the hybrid resulting from the cross between SA49 and CENPK113-1A (SA/CEN) (**Figure 2**).

**Fig 2.**
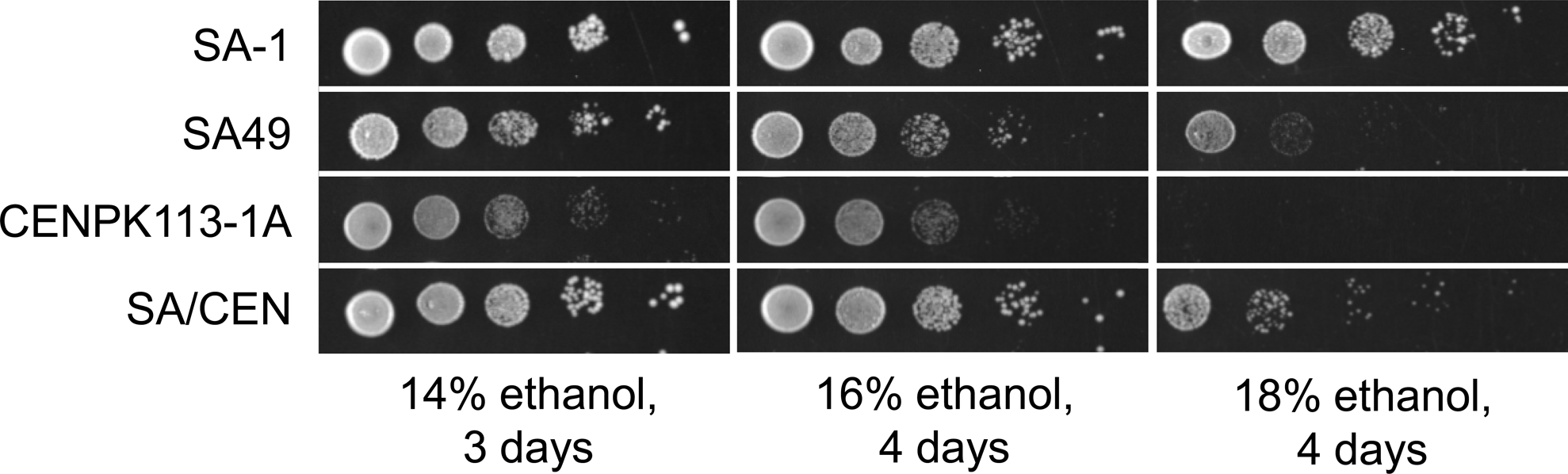
Spot assay in different ethanol concentrations at 30 °C with the parent strains used for QTL mapping.

The spot assay was conducted at higher ethanol concentrations to assess the strains’ response to elevated stress levels. At 14% ethanol, SA-1, SA49, and the SA/CEN hybrid exhibited a similar growth pattern, while no cell growth was observed for the CENPK113-1A strain at the highest dilution. This trend persisted at 16% ethanol, with the laboratory strain experiencing even greater growth inhibition. Remarkably, at 18% ethanol, the industrial strain SA-1 demonstrated exceptional resilience, with minimal impact on cell viability compared to the lowest alcohol concentration. Conversely, haploid SA49 did not maintain its phenotype but still exhibited growth under this condition. The SA/CEN hybrid displayed growth at the lowest dilutions, although cell density was affected. Notably, the laboratory strain CENPK113-1A did not survive at 18% ethanol, confirming its selection for constructing the heterozygous hybrid for QTL mapping.

### QTL mapping

384 segregant haploids from the SA/CEN hybrid were collected and phenotyped in 18% ethanol, the same inhibitory concentration for the susceptible parental strain CENPK113-1A (**Figure 2**). Initially, the entire segregant library was screened under this condition, followed by the selection of the top 95 strains, which were further phenotyped in triplicate. From this assay, two populations of 60 F_2_ segregants, one tolerant and one susceptible to ethanol, were chosen (**Figure S3**). gDNA samples from both pools underwent whole-genome sequencing analysis using the Illumina HiSeq 4000 platform at LaCTAD (Life Sciences Core Facility, Unicamp). Subsequently, the genomic data were analyzed using QTLseqr (Mansfeld and Grumet, 2018) to identify QTL regions associated with the phenotype. A total of 45,410 SNPs were identified between SA49 and CEN.PK113-7D, resulting in the disclosure of 92 potential QTL regions (**Table S3**). The variation of Gprime (G’), the genetic variance attributed to a specific locus, across all chromosomes is depicted in **Figure 3A**. It is evident that numerous loci display elevated G’, suggesting that several genetic markers contribute to the phenotypic variance observed in the population being studied.

**Fig 3.**
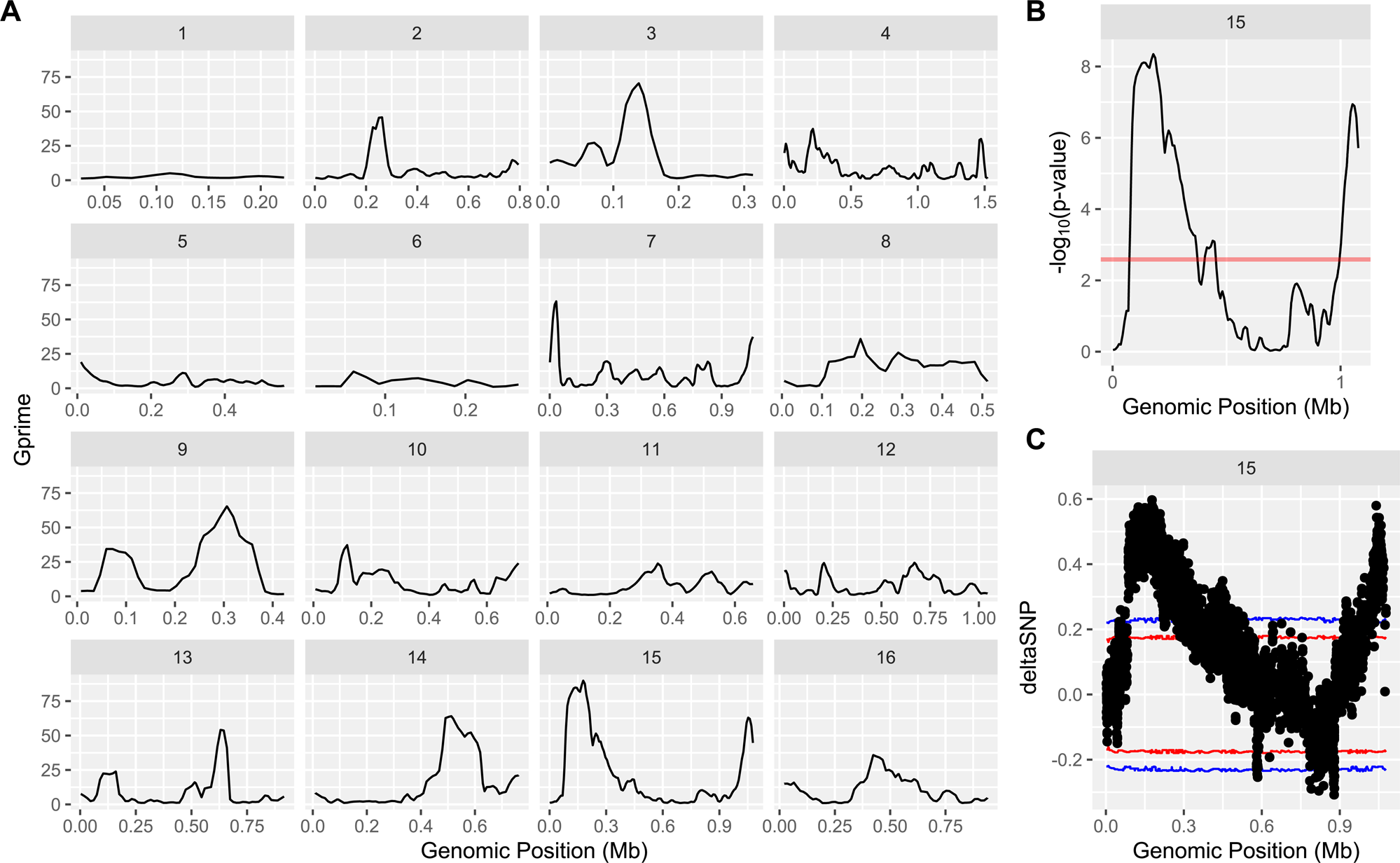
QTL mapping of ethanol resistance in *S. cerevisiae*. A) Variation of Gprime across all 16 chromosomes. B) −log_10_(p-value) in chromosome XV. Red line represents the significant threshold. C) Δ(SNP-index) in chromosome XV. Red and blue dots represent the threshold for a confidence interval of 0.95 and 0.99, respectively.

Due to the intricate nature of ethanol resistance in *S. cerevisiae*, numerous QTL regions were identified in this study, making it challenging to validate genetic targets with significant influence on this trait. Therefore, only regions with a Δ(SNP-index) above a confidence interval (CI) of 0.99 were considered relevant for inferring genotype-phenotype relationships in this study. Following statistical analysis, four QTL peaks were selected for further in-depth examination, located on chromosomes III, VII, XIII, and XV (**Table 2**). Although there are peak regions with negative Δ(SNP-index) values close to the 99% CI, these were not chosen as potential QTL regions due to their association with genomic polymorphisms observed in the low-tolerant segregants pool.

**Table 2:** QTL regions with confidence interval over 0.99 in mapping ethanol resistance in *S. cerevisiae*.

While only four QTL regions met the established threshold as relevant for the studied trait, they still encompass sizable genomic regions, requiring labor-intensive efforts to evaluate the allele contributions to the phenotype. Initially, we decided to delve deeper into the analysis of chromosome XV, as the QTL peak in this region exhibits the lowest p-value and highest Δ(SNP-index) values (**Figure 3B and 3C**). A genomic region spanning 15 kb upstream and downstream from *IRA2* (**Figure 4A**), the gene located at the peak region of this QTL, was selected to identify alleles in SA49 with rare non-synonymous (NS) mutations. These mutations, i.e., alleles with protein mutations not commonly found in other *S. cerevisiae* strains, are more likely to be associated with the ethanol tolerance phenotype. In this genomic interval (chrXV:160000..170000), the only proteins with amino acid changes are Phm7p, Atg34p, Atg19p, Avo1p, Brx1p, and Ira2p (**Table 3**).

**Fig 4.**
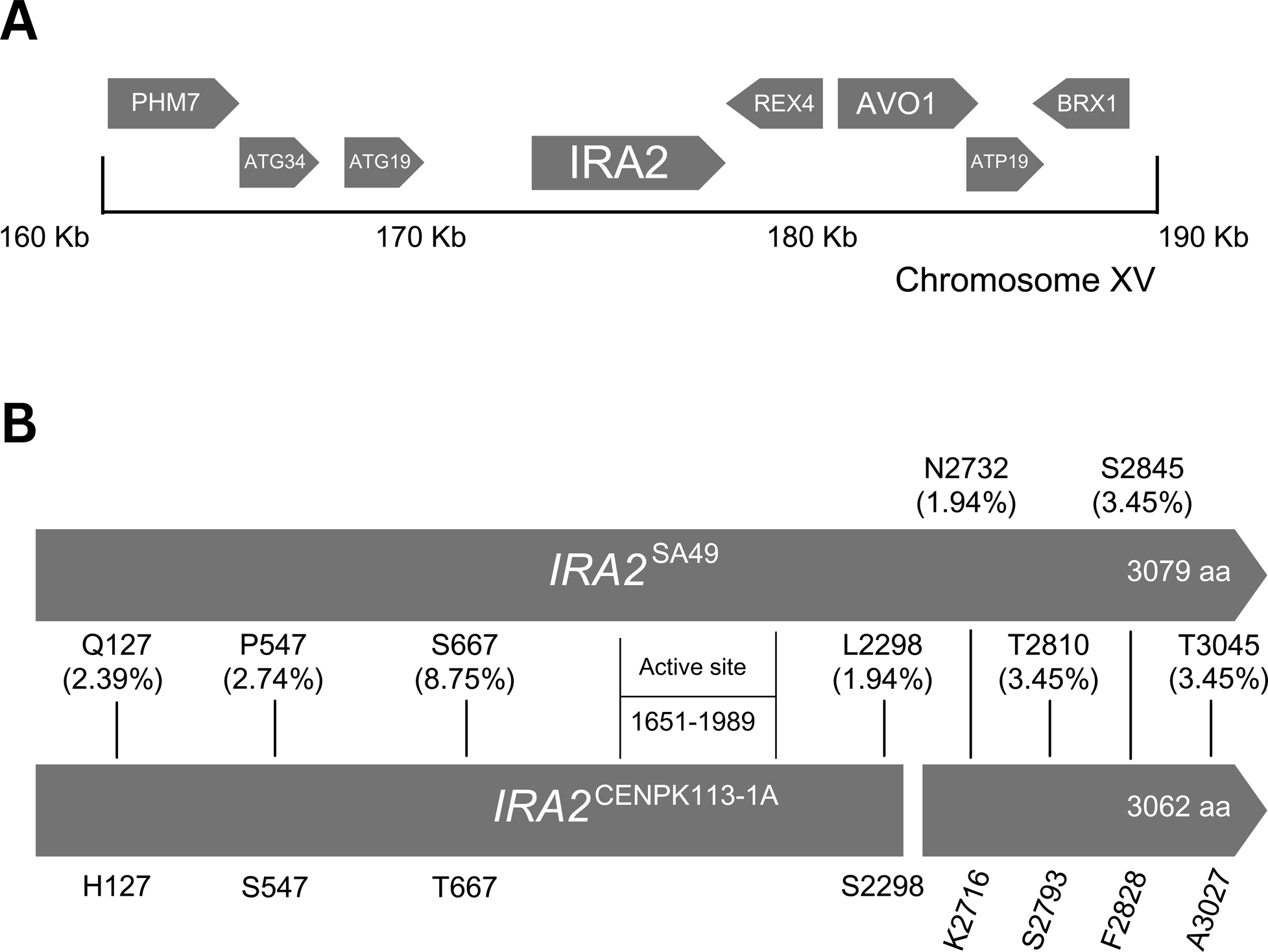
The main genetic architecture studied in this investigation. A) The QTL genomic region validated in this study, spanning 15 kb downstream and upstream *IRA2*, the gene at the peak position on the QTL window in chromosome XV. B) Alignment of *ira2* alleles of SA49 and CENPK113-1A, highlighting the non-synonymous mutations between the alleles and in comparison to a 1011 *S. cerevisiae* population (numbers in parenthesis represent the frequency of the mutations in *IRA2*^SA49^ in the same population).

**Table 3:** Genes with non-synonymous mutations in ethanol-resistant strain SA49 in the interval Chr XV:160000..170000.

From the list of genes with NS mutations, it is evident that *IRA2* exhibits more SNPs than the others. In addition to having more amino acid changes, *IRA2* has been previously reported to be related to other industrially-relevant phenotypes (Parts et al., 2011; Stojiljkovic et al., 2020; Wang et al., 2019), prompting an in-depth analysis of these mutations. To investigate further, the prevalence of such mutations (Q127, P547, S667, L2298, N2732, T2810, S2845, and T3045) in a population of 1011 *S. cerevisiae* strains, previously described (Peter et al., 2018), was calculated. Using the complete sequences of *IRA2* orthologs, the coding sequences were globally aligned, and SNP frequencies were calculated across all strains. SNPs with frequencies lower than 0.1 were considered rare. All eight NS mutations in the SA49 *ira2* allele have frequencies lower than 10%, indicating that these are genetic variants that have been selected in a highly specific environment, considering SA-1 industrial background (Basso et al., 2008; Nagamatsu et al., 2019). Moreover, when compared to the CENPK113-1A allele, the susceptible strain harbors a structural alteration, being shorter (3062 aa) than the SA49 protein (3079 aa) (**Figure 4B**). The larger version of this protein is commonly observed in most strains evaluated. For all the reasons previously exposed, *IRA2*^SA49^ was selected for further molecular validation.

### Reciprocal Hemizygosity Analysis

To assess the influence of *IRA2* on ethanol resistance phenotype, reciprocal hemizygotes were generated. After individually knocking out this allele in the parent strains SA49 and CENPK113-1A, the null mutants were crossed with the other parent expressing the wild-type gene. Two hybrids were created: SA (SA49/CEN.PK113-1A *ira2*Δ::hphMX6), expressing *IRA2*^SA49^; and CEN (SA49 *ira2*Δ::hphMX6/CEN.PK113-1A), expressing *IRA2*^CENPK113-1A^. Both strains, along with the diploid SA/CEN (expressing both alleles) were phenotyped in solid and liquid media under ethanol challenges (**Figure 5**).

**Fig 5.**
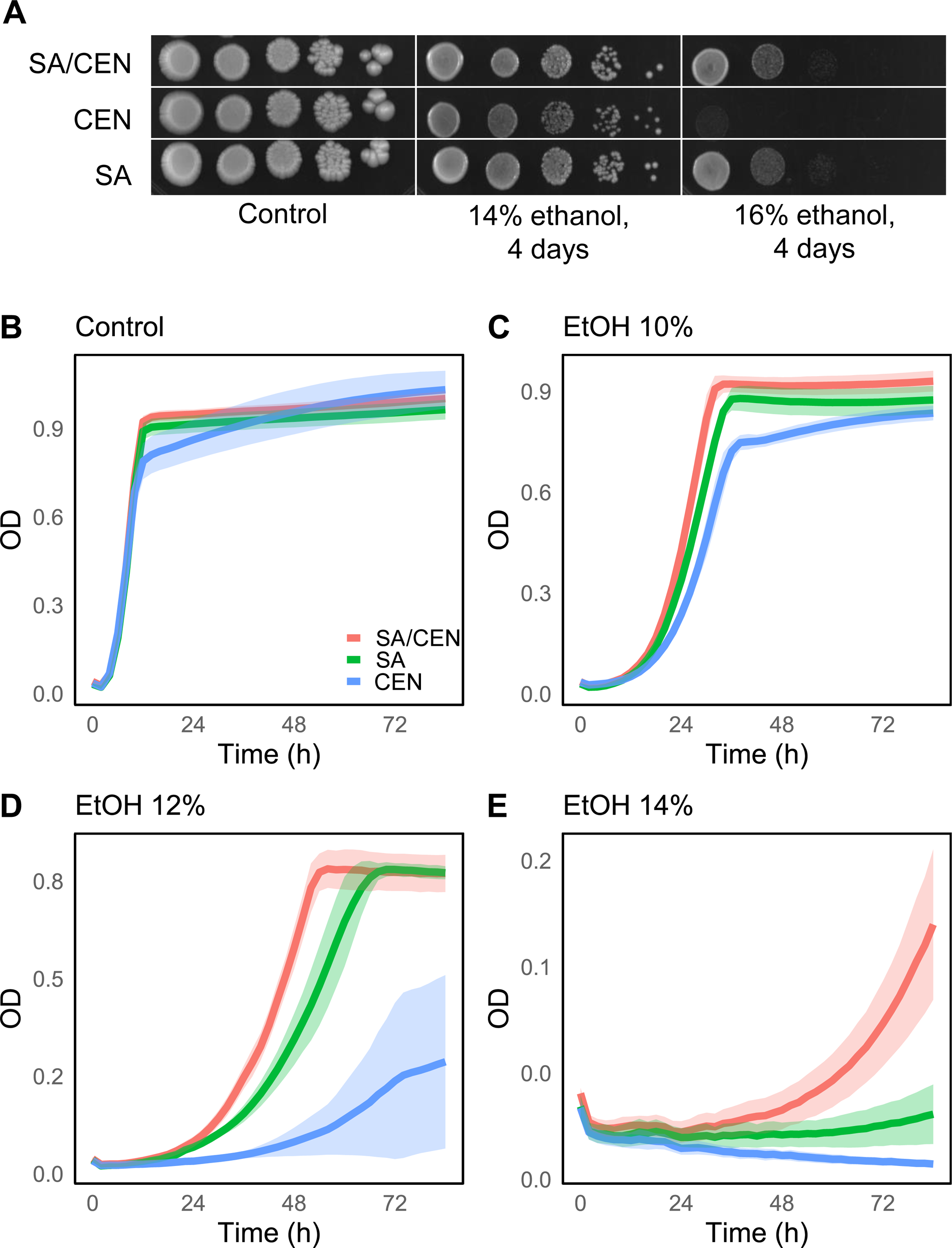
Reciprocal hemizygosity analysis of gene *IRA2* in ethanol stress at 30 °C with strains SA/CEN (expressing both wild-type *ira2* alleles); SA (expressing only *IRA2*^SA49^); and CEN (expressing only *IRA2*^CENPK113-1A^). A) Serial dilutions of tested strains in control (no ethanol), and 14% and 16% ethanol. B) Growth curves of strains in microplate in control, and 10%, 12% and 14% ethanol.

For the results of the spot assay (**Figure 5A**), strains were phenotyped in 14%, 16%, and 18% ethanol final concentrations. Growth of the reciprocal hemizygotes diploids was unaffected under control conditions, suggesting that deletion of one copy of *IRA2* does not impact cell growth. At 14%, both SA and CEN exhibited the same phenotype as strain SA/CEN, with colonies growing even at the lowest dilutions, consistent with previous observations (**Figure 2**). At 16% ethanol, the growth pattern of SA/CEN recapitulated what was previously observed, with a lower cell density. However, at 16% ethanol, the influence of *IRA2* alleles became evident: when the strain expressed only *IRA2*^CENPK113-1A^ (CEN), no cell growth was observed, whereas, in the strain expressing only *IRA2*^SA49^ (SA), the phenotype resembled that of SA/CEN, confirming the influence of the tolerant parent (SA49) allele on ethanol resistance. Although 18% ethanol was tested, no strains were able to grow at this concentration. It is noteworthy that, throughout this study, we noticed some variance between experiments using media with ethanol for phenotyping, possibly due to its volatility, causing subtle variations in final alcohol concentration.

In parallel, the mutant strains were evaluated in liquid media. Initially, testing the same concentrations used for solid media resulted in no growth for all strains at 16% and 18% ethanol, suggesting that ethanol stress in liquid is higher than in solid media due to membrane activity. Therefore, the concentrations of 10%, 12%, and 14% ethanol were used for this assay (**Figure 5B**). The final OD_660_ at 72 hours and μ_MAX_ of all strains in the tested conditions are shown in **Table 4**. When no ethanol is present in the media, no difference in growth pattern is observed between strains SA/CEN, SA, and CEN. At 10% ethanol, there is no difference in the behavior of strains SA/CEN and SA, but the growth of CEN is delayed and its μ_MAX_ lowers. At 12% ethanol, we observe the threshold concentration at which strains phenotype is drastically different: while the hybrid expressing only *IRA2*^SA49^ (SA) is able to reach a final OD_660_ equivalent to the wild-type parent (SA/CEN), the sole expression of *IRA2*^CENPK113-1A^ (CEN) is not sufficient for good performance, and slight growth is only present after 48 hours. In fact, only two CEN replicates, out of four, showed cell growth in this condition. Finally, when using 14% ethanol, after a long adaptation time, SA/CEN reaches a significant OD_660_, and SA presents very little growth. On the other hand, strain CEN had no cell growth at all after 72 hours. Once again, the influence of *IRA2* on the ethanol resistance of yeasts was confirmed.

**Table 4:** Growth parameters of the reciprocal hemizygotes in microplate cultivation at different ethanol concentrations.

### *IRA2* influence in other industrially-relevant phenotypes

As previously mentioned, *IRA2* has been implicated in events related to other stress tolerances, especially due to its role in the negative regulation of the Ras-cAMP signaling pathway, central for cellular response to starvation (Li and Wang, 2013). Therefore, after confirming the influence of this gene in ethanol tolerance, we sought to test if the allele found in SA49 with rare NS mutations is also directly associated with other relevant tolerance traits. Therefore, strains SA/CEN, SA, and CEN were phenotyped at elevated temperatures, low pH, and high acetic acid concentrations (**Figure 6**).

**Fig 6.**
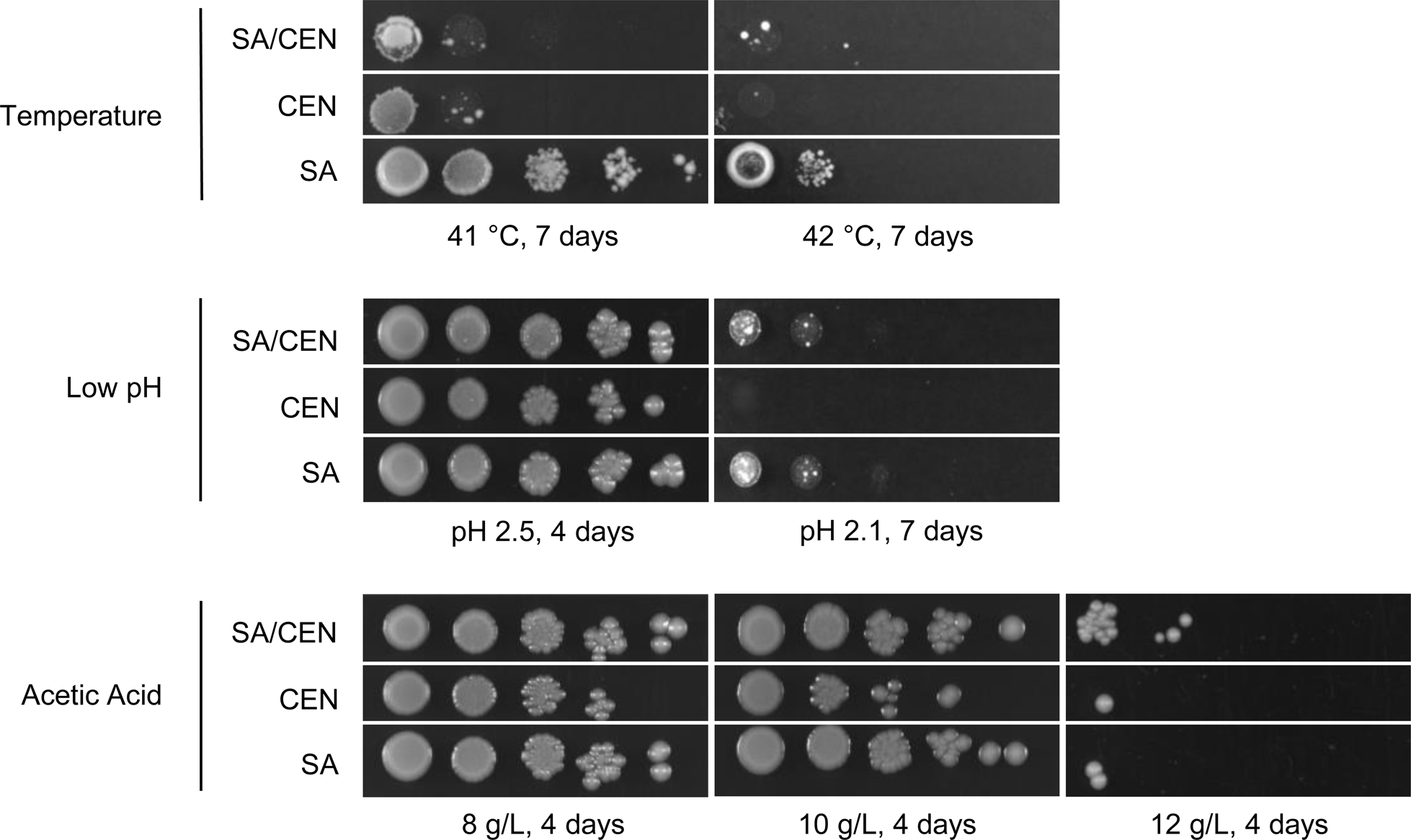
Reciprocal hemizygosity analysis of gene *IRA2* in other industrially relevant stress with strains SA/CEN (expressing both wild-type *ira2* alleles); SA (expressing only *IRA2*^SA49^); and CEN (expressing only *IRA2*^CENPK113-1A^). The yeasts were evaluated in high temperatures (41 and 42 °C), low pH (2.5 and 2.1), and high acetic acid concentrations at pH 4,7 (8, 10, and 12 g/L).

When testing the thermotolerance of the mentioned strains, the effect of *IRA2*^SA49^ was once again positive. At 41 °C, the wild-type hybrid and CEN had similar growth, while SA was able to fully grow even at the lowest cell dilutions. These results suggest a negative effect of *IRA2*^CENPK113-1A^ on thermotolerance. At 42 °C, a highly toxic temperature for yeast, only SA showed some growth. In acidic conditions, pH 2.5 did not hinder growth for any strain tested. On the other hand, at pH 2.1 the growth pattern resembled what happened for ethanol tolerance, where both SA/CEN and SA were able to grow, and the strain expressing only *IRA2*^CENPK113-1A^ (CEN) did not. For acetic acid, at concentrations of 8 and 10 g/L, only CEN growth was hampered, not being able to grow at the lowest dilution. For 12 g/L, another phenomenon was observed, in which only the wild-type diploid had significant growth, and both reciprocal hemizygotes failed to showcase colonies after the second dilution. For this scenario, a dose-dependent response is hypothesized, in which two copies of *IRA2* are necessary for better tolerance towards this organic acid. The strains were also tested in 20 mM 5-hydroxymethyl furfural (HMF) and furfural. After 7 days of cultivation, no differences between cells were observed and, therefore, not shown here.

## DISCUSSION

Several studies have investigated the effect of ethanol on yeast physiology, its molecular tolerance mechanisms, and strategies for enhancing resistance in *S. cerevisiae*. These topics have been recently reviewed (Sahana et al., 2024). The significant interest in exploiting this trait in yeast is due to the crucial role of these microorganisms in various commercial activities, particularly in the brewing and bioethanol industries. In this study, we utilized a simplified QTL mapping approach to identify genetic markers influencing ethanol tolerance in yeast. Our investigation led to the discovery of an *IRA2* allele from an industrial bioethanol strain containing rare NS mutations. In addition to confirming the positive impact of *IRA2* on ethanol tolerance, further analysis revealed its involvement in responses to thermotolerance, low pH, and elevated acetic acid concentrations.

This study began with the selection of a top-performing strain under ethanol challenges, utilizing concentrations of up to 14% alcohol. While *S. cerevisiae* has been documented to thrive in environments containing up to 20% ethanol (Jacobus et al., 2021), a trait typically acquired through adaptive evolution assays, we observed that several wild-type yeasts screened initially did not survive at 14% ethanol. This concentration was therefore chosen as the threshold for ethanol tolerance in subsequent assays. It is noteworthy that while 8% ethanol inhibited the growth of all strains, concentrations exceeding 10% allowed certain strains to produce more biomass than the control condition (with a relative colony size greater than 1.0). Other studies have similarly demonstrated that low ethanol concentrations can enhance cell growth (Sunyer-Figueres et al., 2021). Although other phenomena must be considered, this result can be elucidated by the fact that ethanol can also serve as a carbon source when present at non-toxic concentrations. At last, SA-1 - a well-documented robust industrial bioethanol yeast - was chosen as the parent strain to explore the genetic basis of its ethanol tolerance.

Genetic mapping of the variants associated with the phenotype of interest in strain SA-1 utilized a combined approach of individual phenotyping and bulk whole-genome analysis, followed by a bioinformatics pipeline to identify rare NS mutations in the candidate gene. Previous studies employing a QTL mapping strategy to infer causative genes for ethanol tolerance in yeast have already been documented (Haas et al., 2019; Ho et al., 2021; Riles and Fay, 2019; Swinnen et al., 2012). These investigations have proven the following genes are linked to this phenotype: *MOG1, MGS1*, and *YJR154W* (Haas et al., 2019); *SEC24* (Riles and Fay, 2019); *MKT1*, *SWS2*, and *APJ1* (Swinnen et al., 2012). In our study, 92 potential QTL were uncovered, consistent with prior findings (Haas et al., 2019), affirming that ethanol resistance in *S. cerevisiae* is indeed a complex trait regulated by multiple loci. Assessing all QTL regions and potential causative genes with current technology remains cumbersome, necessitating the prioritization of regions of interest to identify major contributors. Although inbreeding and subsequent selection of large segregant pools for the trait of interest can reduce the size of mapped intervals (Parts et al., 2011), this task remains challenging when evaluating industrially relevant traits. While only four QTL met the established confidence threshold of over 0.99 in this investigation, the regions of interest span large genomic sizes, hindering molecular validation. Therefore, we proposed a filtering strategy that proved highly effective: selecting the QTL with the largest Δ(SNP-index) followed by the analysis of missense mutations within a 15 kb region upstream and downstream of the gene located in the peak position, and finally calculating the frequency of such mutations within an *S. cerevisiae* population.

Within the QTL window selected for in-depth analysis, six genes with NS mutations were identified (*PHM7*, *ATG34*, *ATG19*, *AVO1*, *BRX1*, *IRA2*). Interestingly, polymorphisms in these genes had not been previously linked to ethanol resistance. *AVO1* (which displays 6 NS mutations in SA49) has been documented to be associated with positive effects on high-temperature fermentation (Wang et al., 2019). Conversely, *IRA2* (exhibiting 8 NS mutations in SA49) has not only been correlated with favorable outcomes in high-temperature fermentation (Wang et al., 2019)(REF) but also with acetic acid tolerance (Stojiljkovic et al., 2020) in studies utilizing QTL mapping techniques. Positioned most prominently within the QTL window on chromosome XV and bolstered by previous reports linking this gene to other industrially relevant traits, *IRA2* was chosen to determine the frequency of the NS mutations found in the SA49 allele. Surprisingly, all missense mutations in *IRA2*^SA49^ were deemed rare (with frequencies lower than 0.1 within a population of 1011 strains). Additionally, *IRA2*^CENPK113-1A^ (derived from the susceptible parent strain used in this study) displayed a missing region of 17 amino acids. These factors collectively contributed to the selection of *IRA2* for molecular validation.

The reciprocal hemizygosity analysis for *IRA2* revealed its significant contribution to ethanol resistance in *S. cerevisiae*. On solid media, reciprocal hemizygotes strains exhibited markedly distinct growth patterns at 16% ethanol, with *IRA2*^SA49^ being essential for cell growth at this concentration. Analysis of the phenotype of mutated strains in liquid media yielded noteworthy results at 12% and 14% ethanol concentrations. At these levels, strain SA maintained its performance up to 12%, albeit with a reduced growth rate compared to the parent SA/CEN. However, at 14% ethanol, the absence of *IRA2*^SA49^ in strain CEN led to complete cell death, while the parent and SA strains still exhibited some growth. Given that these ethanol concentrations are commonly found in sugarcane and corn bioethanol mills (Lopes et al., 2016), these results reveal the importance of *IRA2* expression in industrial yeast chassis. While Yoshikawa et al. (2019) demonstrated that *IRA2* deletion hampers cell growth at 8% ethanol using a deletion library (Yoshikawa et al., 2009), our study represents the first documentation of the central role of *IRA2* at elevated ethanol concentration resistance in *S. cerevisiae*.

The investigation of reciprocal hemizygotes under other stresses yielded significant findings. At 41°C, *IRA2*^CENPK113-1A^ impeded cell growth, with only strain SA demonstrating survival. Similarly, at 42°C, strain SA was the sole strain capable of producing colonies. Parts et al. (2011) have previously reported the influence of *IRA2* on thermotolerance using a QTL approach (Parts et al., 2011). Nevertheless, reciprocal hemizygotes were tested only at 40 °C, with a modest difference from the wild-type parent. The authors also suggested that the tested *ira2* allele did not influence other stress responses. Wang et al, (2019) evaluated the *IRA2* influence on fermentation performance in thermal stress of 42 °C, suggesting a minor role of this gene in supporting high temperatures (Wang et al., 2019). Here we present the unequivocal role of *IRA2* tolerance at higher temperatures, as well as a possible negative interaction of an allele (*IRA2*^CENPK113-1A^) to the phenotype and positive influence of the rare NS mutations in *IRA2*^SA49^. At pH 2.1, *IRA2*^SA49^ again proved essential for cell tolerance. Casado et al. (2011) reported the role of *IRA2* at alkaline pH (Casado et al., 2011), but this is the first direct evidence of this gene’s importance to cell growth at inorganic low pH. When 12 g/L acetic acid was introduced to the medium, the SA/CEN hybrid exhibited slightly better performance than the yeast with only one copy of *IRA2*, suggesting a dose-dependent phenotype. The influence of *IRA2* on acetic acid resistance was previously supported (Stojiljkovic et al., 2020).

Further investigation is required to elucidate the genetic alterations underpinning the observed phenotype, whether it be the rare polymorphisms present in SA49 or the 17 amino acids deletion in CENPK113-A Ira2p. Nonetheless, the contribution of *IRA2* is indisputable. Alongside its paralog *IRA1*, *IRA2* functions as a negative regulator of the RAS-cAMP signal transduction pathway by catalyzing the conversion of Ras1,2 from its GTP-active form to the GDP-bound inactive form. Deletion of *IRA2* leads to a constitutively activated Ras-GTP state, resulting in heightened levels of cAMP and protein kinase A (PKA), thereby disrupting the self-regulatory feedback loop of PKAs. This dysregulation of the cAMP-PKA pathway, controlled by *IRA2*, impacts various cellular properties, including thermotolerance, response to starvation, glycogen accumulation, and sporulation capacity (Tanaka et al., 1990; Umebayashi et al., 2001).

As proposed by Stojiljkovic et al. (2020), mutations within *IRA2* may enhance GTPase-stimulating activity, thereby decreasing the activity of the PKA pathway and, ultimately, leading to higher general stress tolerance. Reduced activity of Ras-cAMP/PKA pathway leads to up-regulation of stress response by transcription factors such as Msn2p/Msn4p and Yap1p, which regulates genes involved in ethanol tolerance (Ma and Liu, 2010). Indeed, previous studies have linked polymorphisms in *IRA2* with alterations in growth and stress response (Roop and Brem, 2013) as well as thermotolerance (Wang et al., 2019). Importantly, while there is abundant evidence of *IRA2* on stress tolerance, this gene knockout is related to improved xylose consumption (Myers et al., 2019), a trade-off for second-generation ethanol strains.

In conclusion, the confirmation of *IRA2*’s center role in industrially significant stress responses in *S. cerevisiae* reinforces its importance for biotechnological applications. Simultaneously, it enables the development of robust yeast chassis for commercial applications. Besides, *IRA2*^SA49^ mutations set the stage for elucidating polymorphisms that may enhance Ira2 GTPase-stimulating activity. Moreover, the proposed pipeline for filtering QTL windows to deduce causative alleles enhances the likelihood of pinpointing mutations that underlie a trait of interest. This is particularly valuable considering the challenge of molecularly validating numerous genes, which can impede the identification of major contributors.

## Acknowledgments

The authors thank all the members of our laboratory for their support and scientific advice.

## Author’s contribution

LET, LISS, and BOV performed microbiology and genetic engineering experiments. JJ, MFC, JPG, and EMS performed the in-silico analysis. FSBM, LET and LISS wrote the manuscript. FSBM, MFC, and GAGP supervised this work. FSBM was responsible for the conceptualization of this manuscript. Funding for this work was obtained by GAGP. All authors reviewed the manuscript.

## Data availability

The data presented in this study are available in the article and the complete dataset of DNA-seq reads have been deposited in SRA under accession number PRJNA1109284. Other data are available from the corresponding author upon request.

