## Supplementary material for "Unveiling genetic anchors in Saccharomyces cerevisiae: QTL mapping identifies IRA2 as a key player in ethanol tolerance and beyond": Tables

**Table 1:** Main *S. cerevisiae* strains used in this study.

| Strain | Genotype, description | Reference |
| --- | --- | --- |
| SA-1 | MATa/MATα, bioethanol industrial strain | (Basso et al., 2008) |
| SA49 | MATa, SA-1 ethanol tolerant segregant | This work. |
| CENPK113-1A | MATα, ethanol susceptible laboratory strain | Euroscarf. |
| SA/CEN | MATa/MATα, SA49/CENPK113-1A | This work. |
| SA | MATa/MATα, SA49/CENPK113-1A *ira2*Δ::hphMX6 | This work. |
| CEN | MATa/MATα, SA49 *ira2*Δ::hphMX6/CENPK113-1A | This work. |

**Table 2:** QTL regions with confidence interval over 0.99 in mapping ethanol resistance in *S. cerevisiae*.

| Chromosome | #QTL | Start position (bp) | End position (bp) | Length (bp) | Highest Δ(SNP-index) | Position (bp) | p-value | Gene at peak position |
| --- | --- | --- | --- | --- | --- | --- | --- | --- |
| XV | 1 | 74023 | 372296 | 298273 | 0.5970 | 177492 | 4.68E^-09^ | *IRA2* |
| III | 3 | 101004 | 169637 | 68633 | 0.5478 | 132352 | 6.66E^-08^ | *ADY2* |
| VII | 1 | 1185 | 51986 | 50801 | 0.5284 | 37698 | 2.95E^-07^ | *RAI1* |
| XIII | 3 | 590408 | 666204 | 75796 | 0.5105 | 623104 | 1.20E^-06^ | *CTL1* |

**Table 3:** Genes with non-synonymous mutations in ethanol-resistant strain SA49 in the interval Chr XV:160000..170000.

| Gene | Protein | # Non-synonymous mutations |
| --- | --- | --- |
| *PHM7* | Phm7p | 4 |
| *ATG34* | Atg34p | 6 |
| *ATG19* | Atg19p | 3 |
| *AVO1* | Avo1p | 6 |
| *BRX1* | Brx1p | 1 |
| *IRA2* | Ira2p | 8 |

**Table 4:** Growth parameters of the reciprocal hemizygotes in microplate cultivation at different ethanol concentrations.

| Strain | Control | | 10% Ethanol | | 12% Ethanol | | 14% Ethanol | |
| --- | --- | --- | --- | --- | --- | --- | --- | --- |
|  | OD_600_ | μ_MAX_ (h^-1^) | OD_600_ | μ_MAX_ (h^-1^) | OD_600_ | μ_MAX_ (h^-1^) | OD_600_ | μ_MAX_ (h^-1^) |
| SA/CEN | 0.989 ± 0.020 | 0.575 ± 0.021 | 0.923 ± 0.026 | 0.195 ± 0.008 | 0.773 ± 0.041 | 0.104 ± 0.004 | 0.073 ± 0.017 | 0.044 ± 0.007 |
| SA | 0.954 ± 0.028 | 0.596 ± 0.034 | 0.868 ± 0.036 | 0.184 ± 0.017 | 0.780 ± 0.014 | 0.081 ± 0.012* | 0.025 ± 0.008** | - |
| CEN | 1.016 ± 0.062 | 0.568 ± 0.038 | 0.823 ± 0.019** | 0.152 ± 0.022* | 0.238 ± 0.173** | 0.041 ± 0.023** | 0.008 ± 0.002*** | - |
| (*) p-value < 0.05; (**) p-value < 0.01; (***) p-value < 0.001; (-) not able to calculate | | | | | | | | |
