## Supplementary Information for "Unveiling genetic anchors in Saccharomyces cerevisiae: QTL mapping identifies IRA2 as a key player in ethanol tolerance and beyond"

**Table S1**: Yeast library used in this study for the screening of ethanol tolerance.

| **Strain** | **Species** | **Isolation** | **Origin** |
| --- | --- | --- | --- |
| KAYSER-5 | Saccharomyces cerevisiae (?) | Beer Industry | Department of Genetics ESALQ/USP |
| IZ662 | Saccharomyces cerevisiae | - | Zimotécnico Institute ESALQ/USP |
| IZ2003 | Saccharomyces cerevisiae | - | Zimotécnico Institute ESALQ/USP |
| IZ669 | Saccharomyces cerevisiae | - | Zimotécnico Institute ESALQ/USP |
| M410-A | Saccharomyces cerevisiae | - | Department of Genetics ESALQ/USP |
| IZ1215 | Saccharomyces cerevisiae (?) | Molasses fermentation | Zimotécnico Institute ESALQ/USP |
| IZ1327 | Saccharomyces carsbergensis | - | Zimotécnico Institute ESALQ/USP |
| ATCC26602 | Saccharomyces uvarum | - | American Type Culture Collection - 1982 |
| ATCC28103 | Saccharomyces uvarum | - | American Type Culture Collection - 1982 |
| CAT-1 | Saccharomyces cerevisiae | Sugarcane Biofuel Industry | Basso et al (2008) |
| ANTARTICA-5 | Saccharomyces cerevisiae (?) | Beer Industry | Departamento de Genética ESALQ/USP |
| IZ1169 | Saccharomyces cerevisiae (?) | Wine Industry | Zimotécnico Institute ESALQ/USP |
| IZ1348 | Saccharomyces cerevisiae | - | Zimotécnico Institute ESALQ/USP |
| IZ671 | Saccharomyces cerevisiae | - | Zimotécnico Institute ESALQ/USP |
| M206 | Saccharomyces cerevisiae | - | Department of Genetics ESALQ/USP |
| ATCC4132 | Saccharomyces cerevisiae | Molasses fermentation | American Type Culture Collection - 1982 |
| FLORATIL | Saccharomyces boulardii | - | Departamento de Genética ESALQ/USP |
| IZ237 | Saccharomyces carsbergensis | - | Zimotécnico Institute ESALQ/USP |
| CENPK7D | Saccharomyces cerevisiae | Laboratory | EUROSCARF |
| BG-1 | Saccharomyces cerevisiae | Sugarcane Biofuel Industry | Basso et al (2008) |
| RAD317-8C | Saccharomyces cerevisiae (?) | - | Departamento de Genética ESALQ/USP |
| FT116L | Saccharomyces cerevisiae | Sugarcane Biofuel Industry | Department of Genetics ESALQ/USP |
| IZ1716 | Saccharomyces cerevisisae | - | Zimotécnico Institute ESALQ/USP |
| IZ310 | Saccharomyces cerevisiae | - | Zimotécnico Institute ESALQ/USP |
| X2180-1B | Saccharomyces cerevisiae | - | Department of Genetics ESALQ/USP |
| ATCC26603 | Saccharomyces cerevisiae | - | American Type Culture Collection - 1982 |
| DIAS 164 | Saccharomyces cerevisiae (?) | - | - |
| IZ1099 | Saccharomyces vini | Wine Industry | Zimotécnico Institute ESALQ/USP |
| CENPK122 | Saccharomyces cerevisiae | - | - |
| FM04-58 | Saccharomyces cerevisiae (?) | Sugarcane Biofuel Industry | - |
| FF | Saccharomyces cerevisiae | - | Department of Genetics ESALQ/USP |
| FT119L | Saccharomyces cerevisiae | Sugarcane Biofuel Industry | Department of Genetics ESALQ/USP |
| IZ672 | Saccharomyces cerevisiae | - | Zimotécnico Institute ESALQ/USP |
| IZ659 | Saccharomyces cerevisiae | - | Zimotécnico Institute ESALQ/USP |
| Y10891 | Saccharomyces cerevisiae | Laboratory | EUROSCARF |
| ATCC4125 | Saccharomyces cerevisiae | Molasses fermentation | American Type Culture Collection - 1982 |
| J132B | Saccharomyces diastaticus | - | Labbatt Brewing Co. London Canada - Dpto. Gen. ESALQ |
| ATCC28105 | Saccharomyces uvarum | - | American Type Culture Collection - 1982 |
| JAY270 | Saccharomyces cerevisiae | Sugarcane Biofuel Industry | Department of Genetics ESALQ/USP |
| ETHANOL RED | Saccharomyces cerevisiae | Sugarcane Biofuel Industry | Leaf Lesaffre |
| FMY 097 | Saccharomyces cerevisiae | SA-1 segregant | de Mello et al (2019) |
| IZ677 | Saccharomyces cerevisiae | - | Zimotécnico Institute ESALQ/USP |
| IZ1167 | Saccharomyces cerevisiae (?) | Wine Industry | Zimotécnico Institute ESALQ/USP |
| IZ658 | Saccharomyces cerevisiae | - | Zimotécnico Institute ESALQ/USP |
| Y3575 | Saccharomyces cerevisiae | Laboratory | EUROSCARF |
| ATCC26785 | Saccharomyces cerevisiae | - | American Type Culture Collection - 1982 |
| ATCC28100 | Saccharomyces uvarum | - | American Type Culture Collection - 1982 |
| ATCC28104 | Saccharomyces uvarum | - | American Type Culture Collection - 1982 |
| SA-1 | Saccharomyces cerevisiae | Sugarcane Biofuel Industry | Basso et al (2008) |
| FR | Saccharomyces cerevisiae (?) | Sugarcane Biofuel Industry | - |
| Y1347 | Saccharomyces cerevisiae | - | Department of Genetics ESALQ/USP |
| IZ1351 | Saccharomyces cerevisiae | - | Zimotécnico Institute ESALQ/USP |
| IZ1349 | Saccharomyces cerevisiae | - | Zimotécnico Institute ESALQ/USP |
| IZ651 | Saccharomyces cerevisiae var ellipsoideus | - | Zimotécnico Institute ESALQ/USP |
| Y14620 | Saccharomyces cerevisiae | Laboratory | EUROSCARF |
| IZ137 | Saccharomyces cerevisiae | - | Zimotécnico Institute ESALQ/USP |
| ATCC28097 | Saccharomyces uvarum | - | American Type Culture Collection - 1982 |
| ATCC28102 | Saccharomyces uvarum | - | American Type Culture Collection - 1982 |
| JP-1 | Saccharomyces cerevisiae | Sugarcane Biofuel Industry | - |
| FM04PP | Saccharomyces cerevisiae (?) | Sugarcane Biofuel Industry | Usina coleta por Fox |
| CC-13 | Saccharomyces cerevisiae | - | Department of Genetics ESALQ/USP |
| IZ1350 | Saccharomyces cerevisiae | - | Zimotécnico Institute ESALQ/USP |
| FT117L | Saccharomyces cerevisiae | Sugarcane Biofuel Industry | Department of Genetics ESALQ/USP |
| CAT-1 | Saccharomyces cerevisiae | Sugarcane Biofuel Industry | Basso et al (2008) |
| BG-1 | Saccharomyces cerevisiae | Sugarcane Biofuel Industry | Basso et al (2008) |
| IZ267 | Saccharomyces cerevisiae | - | Zimotécnico Institute ESALQ/USP |
| IZ1827 | Saccharomyces fermentati | - | Zimotécnico Institute ESALQ/USP |
| ATCC28099 | Saccharomyces uvarum | - | American Type Culture Collection - 1982 |
| BY4742 | Saccharomyces cerevisiae | Laboratory | - |
| SCHIN-5 | Saccharomyces cerevisiae (?) | Beer Industry | Departamento de Genética ESALQ/USP |
| IZ2004 | Saccharomyces cerevisiae | - | Zimotécnico Institute ESALQ/USP |
| IZ1832 | Saccharomyces cerevisiae | - | Zimotécnico Institute ESALQ/USP |
| M4-10B | Saccharomyces cerevisiae | - | Department of Genetics ESALQ/USP |
| ATCC24858 | Saccharomyces cerevisiae | - | American Type Culture Collection - 1982 |
| IZ287 | Saccharomyces cerevisiae var ellipsoideus | - | Zimotécnico Institute ESALQ/USP |
| IZ1904 | Saccharomyces boulardii | - | Zimotécnico Institute ESALQ/USP |
| J1741C | Saccharomyces diastaticus | - | Labbatt Brewing Co. London Canada - Dpto. Gen. ESALQ |
| BY4741 | Saccharomyces cerevisiae | Laboratory | EUROSCARF |
| (?) means the species confirmation was not clear. Missing information are identified with (-). | | | |

**Table S2**: Primers used in this study.

| **Label** | **Sequence (5’-3’)** | **Description** |
| --- | --- | --- |
| LTO_001 | AACATTTAACCACATTTTAGCACAC | *IRA2* amplification |
| LTO_002 | CGATAGAATATATAAAAATTGTAGCTGTATAAAA | *IRA2* amplification |
| LTO_003 | TTCGGATAACGAAAATGCAA | *IRA2* internal |
| LTO_004 | TCAACTAAACTGTATACATTATCTTTCTTCAGGGAGAAGCGCGCCAGATCTGTTTAGCTT | hphMX6 amplification from pAG32 |
| LTO_005 | ATAGATATTGATATTTCTTTCATTAGTTTATGTAACACCTGCCGATTCATTAATGCAGGT | hphMX6 amplification from pAG32 |
| LTO_006 | gatcAAGCTGAAAAGATCAAAGTG | sgRNA* to be cloned in pGS |
| LTO_007 | aaacCACTTTGATCTTTTCAGCTT | sgRNA* to be cloned in pGS |
| *sgRNA sequence in uppercase | | |

**Table S3**: All QTL regions disclosed throughout the chromosomes.

| **CHROM** | **qtl** | **start** | **end** | **length** | **nSNPs** |
| --- | --- | --- | --- | --- | --- |
| II | 1 | 204754 | 283795 | 79041 | 332 |
| II | 2 | 771152 | 771202 | 50 | 2 |
| III | 1 | 14287 | 15554 | 1267 | 8 |
| III | 2 | 49664 | 83170 | 33506 | 140 |
| III | 3 | 101004 | 169637 | 68633 | 171 |
| IV | 1 | 2828 | 34775 | 31947 | 117 |
| IV | 2 | 167905 | 344825 | 176920 | 727 |
| IV | 3 | 379282 | 394389 | 15107 | 90 |
| IV | 4 | 1457135 | 1495071 | 37936 | 241 |
| V | 1 | 9994 | 21005 | 11011 | 24 |
| VII | 1 | 1185 | 51986 | 50801 | 196 |
| VII | 2 | 277647 | 315685 | 38038 | 80 |
| VII | 3 | 574961 | 579753 | 4792 | 41 |
| VII | 4 | 767077 | 789078 | 22001 | 63 |
| VII | 5 | 826892 | 841935 | 15043 | 26 |
| VII | 6 | 1026949 | 1063570 | 36621 | 102 |
| VIII | 1 | 112569 | 241750 | 129181 | 464 |
| VIII | 2 | 263280 | 488145 | 224865 | 613 |
| IX | 1 | 46254 | 123718 | 77464 | 390 |
| IX | 2 | 224814 | 373797 | 148983 | 422 |
| X | 1 | 87083 | 135765 | 48682 | 196 |
| X | 2 | 162069 | 268589 | 106520 | 654 |
| X | 3 | 678811 | 709493 | 30682 | 93 |
| XI | 1 | 301663 | 371799 | 70136 | 251 |
| XI | 2 | 497131 | 535576 | 38445 | 164 |
| XII | 1 | 1706 | 18063 | 16357 | 69 |
| XII | 2 | 186529 | 234030 | 47501 | 253 |
| XII | 3 | 611618 | 617965 | 6347 | 23 |
| XII | 4 | 626773 | 716060 | 89287 | 321 |
| XII | 5 | 761791 | 778040 | 16249 | 59 |
| XIII | 1 | 85586 | 162358 | 76772 | 345 |
| XIII | 2 | 507221 | 522116 | 14895 | 24 |
| XIII | 3 | 590408 | 666204 | 75796 | 222 |
| XIV | 1 | 417972 | 629188 | 211216 | 613 |
| XIV | 2 | 727047 | 761131 | 34084 | 232 |
| XV | 1 | 74023 | 372296 | 298273 | 1432 |
| XV | 2 | 403638 | 452863 | 49225 | 275 |
| XV | 3 | 995622 | 1077579 | 81957 | 322 |
| XVI | 1 | 21771 | 44878 | 23107 | 146 |
| XVI | 2 | 53816 | 70505 | 16689 | 23 |
| XVI | 3 | 357396 | 623248 | 265852 | 719 |

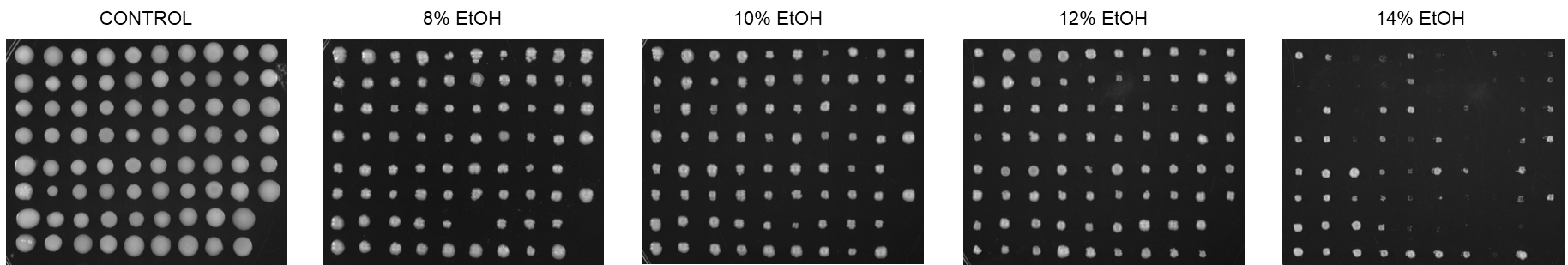

**Figure S1**: Screening of 78 yeast strains in different ethanol concentrations at 96 hours.

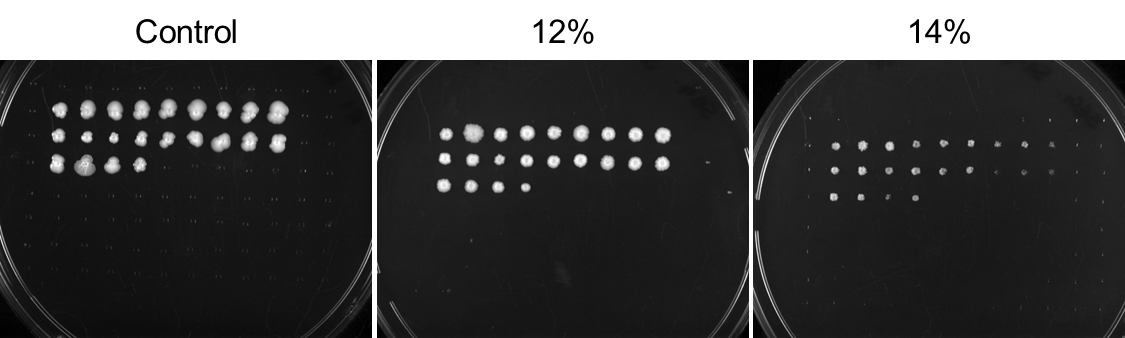

**Figure S2**: SA-1 segregants screening in different ethanol concentrations at 96 hours.

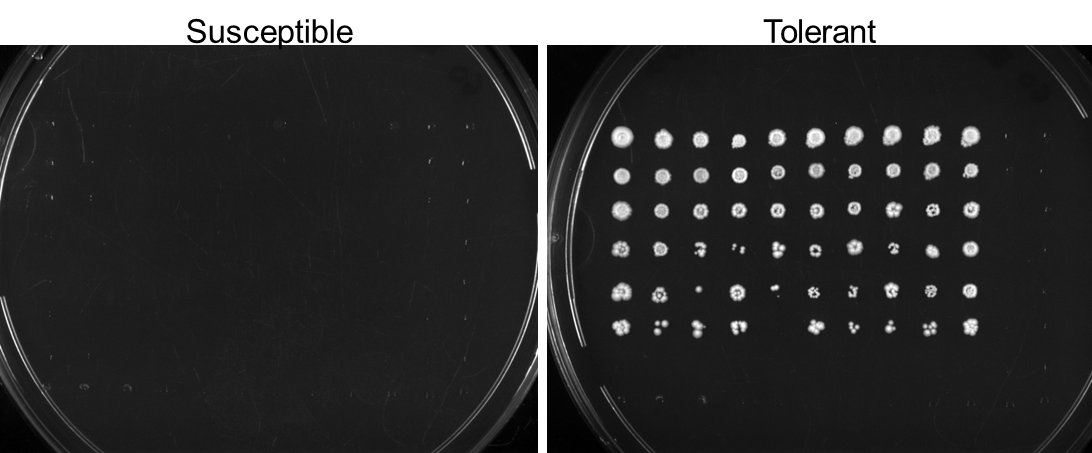

**Figure S3**: Phenotyping of the two 60 individuals population for QTL mapping.
